# Machine learning approaches based on fibroblast morphometry confidently identify stress but have limited ability to predict ALS

**DOI:** 10.1101/2022.10.23.513410

**Authors:** Csaba Konrad, Evan Woo, Kirsten Bredvik, Bangyan Liu, Thomas J. Fuchs, Giovanni Manfredi

**Affiliations:** Brain and Mind Research Institute, Weill Cornell Medicine, New York, NY, USA; Hasso Plattner Institute for Digital Health, Icahn School of Medicine at Mount Sinai

**Keywords:** Amyotrophic lateral sclerosis, Fibroblasts, Machine learning, Microscopy, Biomarkers

## Abstract

**Objective:** Amyotrophic lateral sclerosis (ALS) is a devastating neuromuscular disease with limited therapeutic options. Diagnostic and surrogate endpoint biomarkers are needed for early disease detection, clinical trial design, and personalized medicine.

**Methods:** We tested the predictive power of a large set of primary skin fibroblast (n=443) from sporadic and familial ALS patients and healthy controls. We measured morphometric features of endoplasmic reticulum, mitochondria, and lysosomes by imaging with vital dyes. We also analysed immunofluorescence images of ALS-linked proteins, including TDP-43 and stress granule components. We studied fibroblasts under basal conditions and under metabolic (galactose medium), oxidative (arsenite), and heat stress conditions. We then employed machine learning (ML) techniques on the dataset to develop biomarkers.

**Results:** Stress perturbations caused robust changes in the measured features, such as organellar morphology, stress granule formation, and TDP-43 mislocalization. ML approaches were able to predict the perturbation with near perfect performance (ROC-AUC > 0.99). However, when trying to predict disease state or disease groups (e.g., sporadic, or familial ALS), the performance of the ML algorithm was more modest (ROC-AUC Control vs ALS = 0.63). We also detected modest but significant scores when predicting clinical features, such as age of onset (ROC-AUC late vs early = 0.60).

**Conclusions:** Our findings indicate that the ML morphometry we developed can accurately predict if human fibroblasts are under stress, but the differences between ALS and controls, while statistically significant, are small and pose a challenge for the development of biomarkers for clinical use by these approaches.

## Introduction

Amyotrophic lateral sclerosis (ALS) is a devastating neurodegenerative disease, characterized by rapidly progressive loss of upper and lower motor neurons, resulting in paralysis and death within 3-4 years after disease onset(1, 2). Approximately 90% of cases are sporadic (sALS) with no known family history of the disease; however, the genetic component of developing ALS is estimated between 21-65%(3-7). While progressive motor neuron loss, decline of motor function, and TDP-43 aggregation are the most common denominators, the clinical presentation, such as progression rate, onset location, age at onset, and presence of co-morbidities, is variable among patients(8). Many processes are involved in ALS pathogenesis, including RNA metabolism, protein homeostasis, nuclear transport, excitotoxicity, mitochondrial dysfunction, oxidative stress, stress granule dynamics, axonal transport, neuroinflammation, glial dysfunction, and vesicular transport defects(2, 9-11).

The broad diversity of molecular mechanisms contributing to the disease could explain the observed clinical complexity, implying that the ability to define subgroups with shared disease mechanisms is of utmost importance for developing effective therapies for ALS. Therefore, the search for biomarkers that predict clinical features, define subgroups, or point towards new therapeutic targets is a high priority in the field of ALS research, especially for the sporadic forms. Recent efforts have yielded several promising biomarkers based on biofluids (blood, CSF, saliva, urine), biopsies (nerve, skin, muscle), patient cells (fibroblasts, PBMCs), and imaging (cerebral MRI, spinal MRI, PET), however, they are all in their early validation phase, yet to reach the clinic.

Several lines of evidence point to differences in sALS skin compared to controls, such as increased collagen density(12, 13), increased MMP-2 and MMP-9(14, 15), increased S100A4(16), and an altered aging metabolism phenotype(17). Furthermore, ALS skin and fibroblasts have been shown to recapitulate pathology thought to play key roles in neuronal death, such as SOD1 misfolding(18), FUS(19) and TDP-43(20-27) pathology, mitochondrial dysfunction(28-30), abnormal stress granule formation(31, 32) and RNA processing(33), as well as altered protein homeostasis in the case of C9Orf72 mutant fibroblasts(34).

We hypothesized that some cellular processes involved in MN death in sALS may also be present in patient-derived skin fibroblasts. Previously, we found higher mitochondrial membrane potential and elevated rates of oxidative and glycolytic metabolism in patient derived fibroblasts from sporadic ALS, primary lateral sclerosis (PLS)(35), and C9Orf72 ALS patients(36, 37). In a follow up study, we confirmed the hypermetabolic phenotype of sALS fibroblasts and identified a subgroup with accelerated metabolic flux through their transsulfuration pathway that we suspect serves to increase glutathione production as a defence against oxidative stress(38). These findings support the use of fibroblasts for studying the pathogenic mechanisms of ALS, although metabolic biomarkers need to be further developed in order to be applied to the clinic. In parallel to metabolism, readouts involving microscopy are being explored for fibroblast-based ALS biomarker development(21-23, 25, 26, 31), although most of these studies are on small numbers of sALS cell lines (except(22)), and one study reported negative results(39).

In this study, taking advantage of a large number (n=443) of de-identified human fibroblast lines available in our repository, we tested if fibroblast morphometry can be used as a biomarker for ALS. We first focused on live cell imaging of key organelles, including mitochondria, endoplasmic reticulum (ER), and lysosomes. We then performed immunocytochemistry for proteins known to be involved in ALS. Taking advantage of the option to subject live cells to stress, we imaged all lines under baseline and stress perturbations, such as forced oxidative metabolism in galactose medium, oxidative stress by arsenite, and temperature stress. Through this effort, we generated an annotated dataset suitable for learning tasks by ML. Using this strategy, we found robust changes in cell morphometry caused by stress perturbations that make them easily recognizable using ML approaches, and there were subtle differences between ALS and controls that could be used to build models predictive of ALS with limited but significant performance.

## Materials and Methods

### Skin Biopsy, fibroblast cultures and stress perturbations

After informed consent, a punch skin biopsy was obtained from the volar part of the forearm. Skin biopsies were de-identified to protect patients’ identity. Fibroblast samples were provided to our laboratory as coded samples. Some lines were obtained from the NINDS catalogue of motor neuron disease fibroblasts. Skin fibroblasts were cultured as described previously(36-38) in Dulbecco modified Eagle medium (DMEM) (ThermoFisher Scientific, Waltham, MA) supplemented with 25 mM glucose, 4 mM glutamine, 1 mM pyruvate, and 10% foetal bovine serum (hereafter growth medium). All cultured fibroblast lines were studied at passages ranging between 5 and 12. We have not observed loss of contact inhibition in any of the lines.

Fibroblast lines were replated in 96 well glass-bottom imaging plates (P96-1.5H-N, Cellvis) at a density of 8000 cells/well in experimental media (DMEM (A1443001), 10% dialyzed FBS, Ab+F, 2.5 ug/ml plasmocin, 2 mM glutamax, 1 mM pyruvate) supplemented with either 5 mM glucose or 5 mM galactose.

The next day we subjected all fibroblast patient lines to 6 different stress conditions: 1) baseline: no stress, 2) galactose: as described above, 3) arsenite: to cause oxidative stress 1mM arsenite was added to the medium for 30 minutes, 4) arsenite post stress: the medium was replaced with experimental media, and cells were left to rest for 90 minutes, 5) heat shock: cells were placed in an incubator set to 44ºC for 1 hour 6) heat shock post stress: after the heat shock cells were left to rest for 90 minutes at 37 ºC.

### Live imaging (LI)

Cells were seeded in 96 well imaging plates and subjected to stress perturbations (see above). Fibroblasts were then loaded with Hoechst 33342 (1 µg/ml), ER tracker green (1 µM), TMRM (15 nM) and Lysotracker deep red (50 nM) for 30 minutes. Images (20 fields) were taken on an ImageXpress pico (Molecular devices) at 40x magnification (pixel size = 0.1725 µm).

### Immunofluorescence (IF)

Cells were seeded and treated identically as above, but wells were fixed for 30 minutes in 4% paraformaldehyde, blocked by BSA in PBS+20 mM glycine, consequently incubated over night with anti TDP-43 rabbit (Proteintech, cat 10782-2-AP, 1:200), anti G3BP1 mouse (Santa cruz, cat sc-81940, 1:200) and anti HSP60 (Enzo, cat: ADI-SPA-828-F, 1:100) at 4 ºC, followed by washing and staining by Cy2 anti-goat, Cy3 anti-mouse, Cy5 anti-rabbit (Jackson, 1:500), and Hoechst 33342 (1 µg/ml). Images (30 fields) were taken as described above.

### Hand engineered automated feature extraction

As shown on Fig. 2 the four fluorescent channels were used in both imaging modalities. First, background was subtracted in all channels (r=500px=86.25µm). Cell nuclei were segmented by gaussian blur (r=15px=2.59µm), adaptive thresholding and watersheding. To get cell bodies, we first pixelwise summed the four channels, blurred and thresholded, then fed the image with the segmented nuclei as seeds into a Voronoi algorithm using Cellprofiler. Objects on the edge were discarded and single cell measurements were carried out in ImageJ. When ultrastructural segmentation was needed (mitochondria, lysosomes, TDP-43, G3BP1, HSP60), the individual cells were cropped and segmented similarly as the nuclei, but with appropriate kernel sizes for blur and size exclusion. Circular objects (lysosomes, G3BP1 and TDP-43 granules) were watersheded, while mitochondrial stains were not and were subjected to skeletonization and analysis. Organellar level data was then integrated into cell level data and consequently into patient level data in Python.

**Figure 1.**
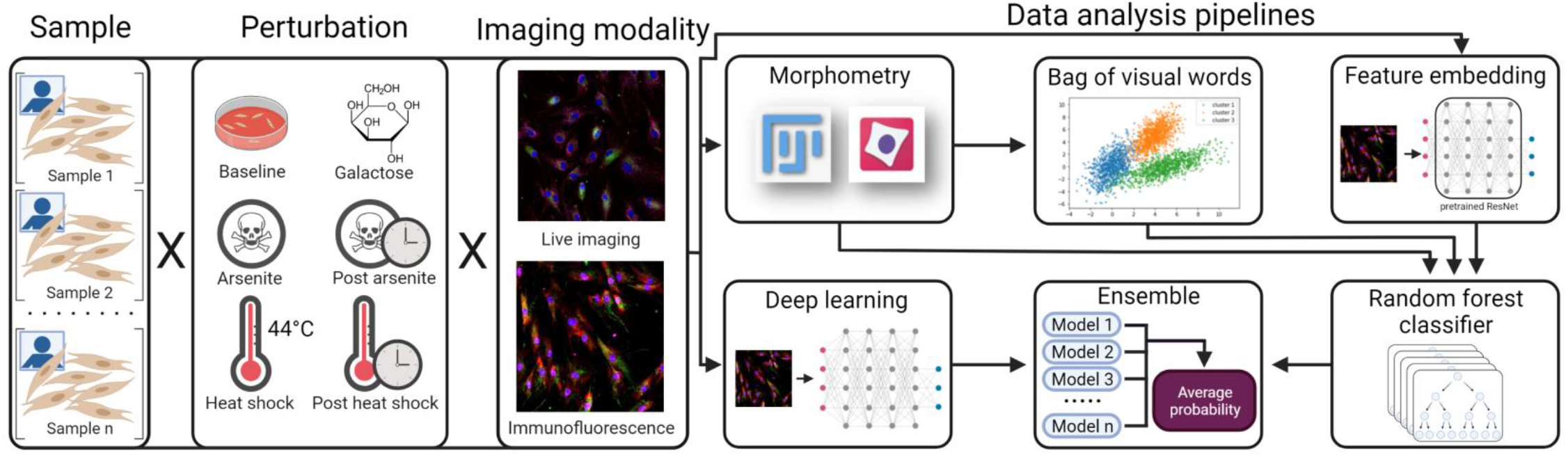
Flow chart of data collection and analysis pipelines. Patient and control primary skin fibroblasts were subjected to five stress perturbations: 1) culturing in galactose medium overnight, 2) 30 min incubation with of 1mM arsenite, 3) 30 min incubation with arsenite + 90 min of recovery time, 4) 60 min of heat shock at 44°C, 5) 60 min of heat shock + 90 min of recovery time. Cells were live stained with dyes recruited to specific organelles (ER tracker, TMRM, Lyso tracker) or fixed and immunostained for TDP-43, G3BP1, and HSP60. Images were taken with a high content imager (at 40X magnification) and processed by scripts extracting specific morphometric measurements. Four machine learning approaches (morphometry + random forest, bag of visual words + random forest, feature embedding + random forest, deep learning from scratch) were used to make predictions.

**Figure 2.**
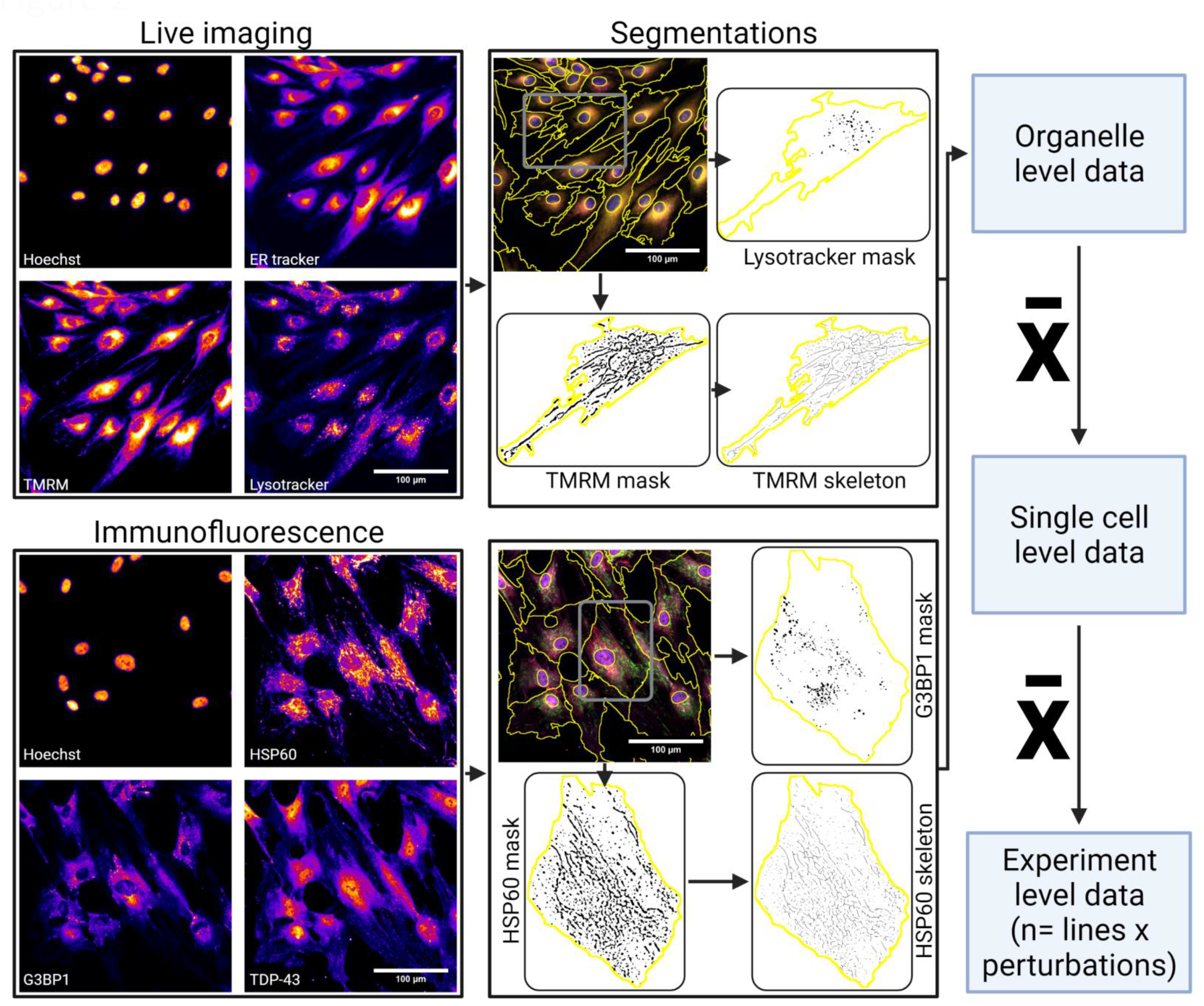
Flow chart of automated morphometric feature extraction. Specific scripts were written in Fiji ImageJ and Cellprofiler to batch process the images. Single fibroblasts were first identified by their nuclei and then the cell body was segmented by using the 3 other fluorescent channels. Individual cells were cropped and measurements performed (as described in Table S1). The data was then integrated from organellar level to cell level and lastly at the patient cell line level.

### Statistical analyses

We used Two-way ANOVA (independent variables: stress perturbation, disease group) implemented Python (statsmodels.stats.anova.anova_lm), with p-value correction for multiple comparisons using the Benjamini-Hochberg method, with false discovery rate set to 0.05. For the features that passed, we performed Tukey’s ‘Honest Significant Difference’ method in both directions of independent variables.

### Random forest optimization

For the three ML approaches utilizing random forests, the sklearn.ensemble. RandomForestClassifier library was used in python. Model parameters (number of estimators, max features, max depth, min samples/split, min samples/leaf, class weighting) were optimized using random grid search on the training set with 3-fold cross-validation for 300 iterations. After getting the best parameters, 10-fold crossvalidation was used to get the cross-validation metrics. Following cross-validation, the model was trained on all the training data and tested on the test set.

### Bag of Visual Words

For the BOVW approach, k-means clustering is used to cluster the cells and generate histograms that are used as features by the RF algorithm. We determined the ideal number of clusters by testing clustering from k = 5-300, training a random forest on the data to predict stress perturbation and measuring ROC-AUC and accuracy metrics for the cross-validation and test sets. We saw no significant improvement in either imaging modalities above k=25 (Fig. S3A-B), and therefore 25 clusters were used to generate the histograms features for all classification tasks reported in Tables 2 and S5.

### Feature embedding and deep learning

Feature embedding and deep learning were done similar to(40) using the pytorch deep learning library, and for both we used a ResNet 50 architecture. For both, 244×244 pixel images were cropped around 100 cells using nuclei centroid coordinates from the morphometric pipeline. For feature embedding, weights learned on ImageNet were used, the 5 channels were passed through the network seperately (triplicating each to match the input dimensions of the network), and the last linear layer activations (2048 features) were used. Each channel was then compressed from 2048 to 64 features using PCA to produce a final vector of size 320 for the 5 channels for each cell image. 64 features was chosen by measuring variance retained by PCA. We found that at 64 features 65-80% variance was retained (Fig. S3C). The 320 features were then averaged for the 100 cells to get to patient level data for each stress perturbation, and RF was used for the classification tasks. For deep learning from sratch, the Resnet architecture was modified to accommodate 5 input channels and 6 or 2 outputs, depending on the task. The same cropped 224×224 pixel cells images were used, with 5x cross-validation for 10 epochs with learning rate decay (halving each epoch) and stochastic gradient descent optimization. For each cell image instead of the predictions (logits), class probabilities were recorded and averaged to get a final prediction. An example of training curves is shown on Fig. S3D (IF stress perturbation prediction, training on all training examples, testing on test set, measured on the cell level).

## Results

### Stress perturbations have robust and reproducible effects on fibroblast morphometry regardless of disease state

In this study we used two imaging modalities: 1) simultaneous live imaging (LI) with vital dyes of the endoplasmic reticulum, mitochondria, and lysosomes; 2) simultaneous immunofluorescence (IF) of TAR DNA-binding protein 43 (TDP-43), the stress granule assembly component Ras GTPase-activating protein-binding protein 1 (G3BP1), and the mitochondrial chaperone heat-shock protein 60 (HSP60). These proteins were selected based on their proposed roles in ALS pathogenesis, and because they have been found to be altered in ALS fibroblasts(41-43). In both imaging modalities, in addition to these three channels, nuclei were stained with a vital dye and brightfield images collected, resulting in a total 5 channels per modality. Table S1 lists the morphometric features measured in the two imaging modalities.

We analysed a total of 137 healthy control, 225 sALS, 41 primary lateral sclerosis (PLS), 26 C9Orf72 mutant, and 14 SOD1 mutant fibroblast lines. A morphometry pipeline was developed using ImageJ and CellProfiler to take the measurements we deemed relevant (hand engineered features, Fig. 2). The measurements were taken at the organellar level (e.g., mitochondrial length or lysosomal intensity) and integrated at the cell level features by averaging or summation (e.g., number of mitochondria or G3BP1 granule per cell), and then integrated to subject level data by averaging the cell level data (Fig. 2). Fig. S1 list all the comparisons of morphometric features measured in the two imaging modalities (LI and IF) under the different stress conditions.

First, we assessed the efficacy of the stress perturbation on altering morphometric parameters in all cell lines, independent of disease state (i.e., ALS or control). Subjecting fibroblasts to stress perturbations resulted in abundant morphometric alterations. There were statistically significant differences between culture conditions for all 55 features extracted from LI and IF (Fig. S1). Specific perturbations, however, had distinct effects on the features. For example, all perturbations caused a significant decline in mitochondrial membrane potential measured as TMRM signal (Fig. 3A), but only after arsenite oxidative stress, mitochondria recovered membrane potential and became hyperpolarized. This finding suggests that cellular energy demands are different after oxidative stress than after heat stress. Another interesting observation was that lysotracker intensity, indicative of the acidity of lysosomes, declined immediately upon arsenite stress and further decreased post stress, while it only declined post stress upon heat shock. This suggests that the effects of oxidative stress on lysosomal ATPase are more acute than heat shock. Similarly, stress granules formed immediately after arsenite stress but only post stress in the case of heat shock. On the contrary, TDP-43 mislocalized from nucleus to cytosol immediately after heat shock but only after post stress for arsenite. Interestingly, a lack of TDP-43 mislocalization was also reported in earlier studies, where fibroblasts were not imaged post arsenite stress(32, 39).

**Figure 3.**
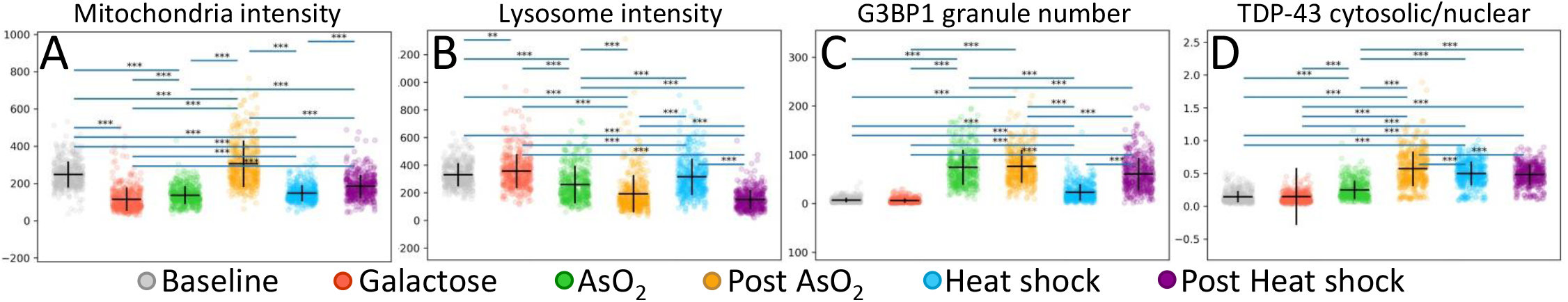
Examples of four morphometric changes caused by stress perturbations (mitochondrial initensity, lysosome intensity, G3BP1 granule number, TDP-43 cytosolic to nuclear intensity ratio). Detailed description of features and measurement units can be found in Table S1. The complete lists of significant changes is shown in Fig. S1. Each point represents a fibroblast line (n=443); horizontal lines represent the mean; vertical lines represent the standard deviation; *: p-adj<0.05; **: p-adj<0.01; ***: p-adj<0.001.

Overall, the comparative analyses of LI and IF features before, during, and after stress condition, provide evidence that the imaging pipeline we developed is adequate to detect cellular changes caused by various types of stress.

### ALS fibroblasts show modest differences between groups

We next compared morphometric features of control, sALS, PLS, C9Orf72 and SOD1 mutant fibroblasts, independent of perturbation. We found differences between groups with small effect sizes. Fig. S2 shows all comparisons between groups that passed statistical significance test. Notably, both HSP60 and TMRM staining of mitochondria showed that sALS fibroblasts have less mitochondria per cell than controls and SOD1 fibroblasts (Fig. 4A, D). SOD1 cells also displayed a more complex network of HSP60 objects compared to other ALS groups and controls. The number of lysosomes per cell was smaller in sALS fibroblasts compared to controls (Fig. 4B). On the other hand, sALS fibroblasts had higher G3BP1-derived features, such as granule area (Fig. 4E), granule diameter, granule intensity and total cellular G3BP1 intensity (Fig. S2). Moreover, some TDP-43 related features were higher identified in PLS fibroblasts than in other groups. The colocalization of TDP-43 with G3BP1 was increased, as well as total cellular TDP-43 staining, compared to sALS. The intensity of segmented TDP-43 granules was also higher compared to controls and sALS. Furthermore, we found that SOD1 fibroblasts exhibit similar nuclear sizes but more elliptically shaped nuclei, as evidenced by the difference of the feret diameter, compared to all other ALS groups.

**Figure 4.**
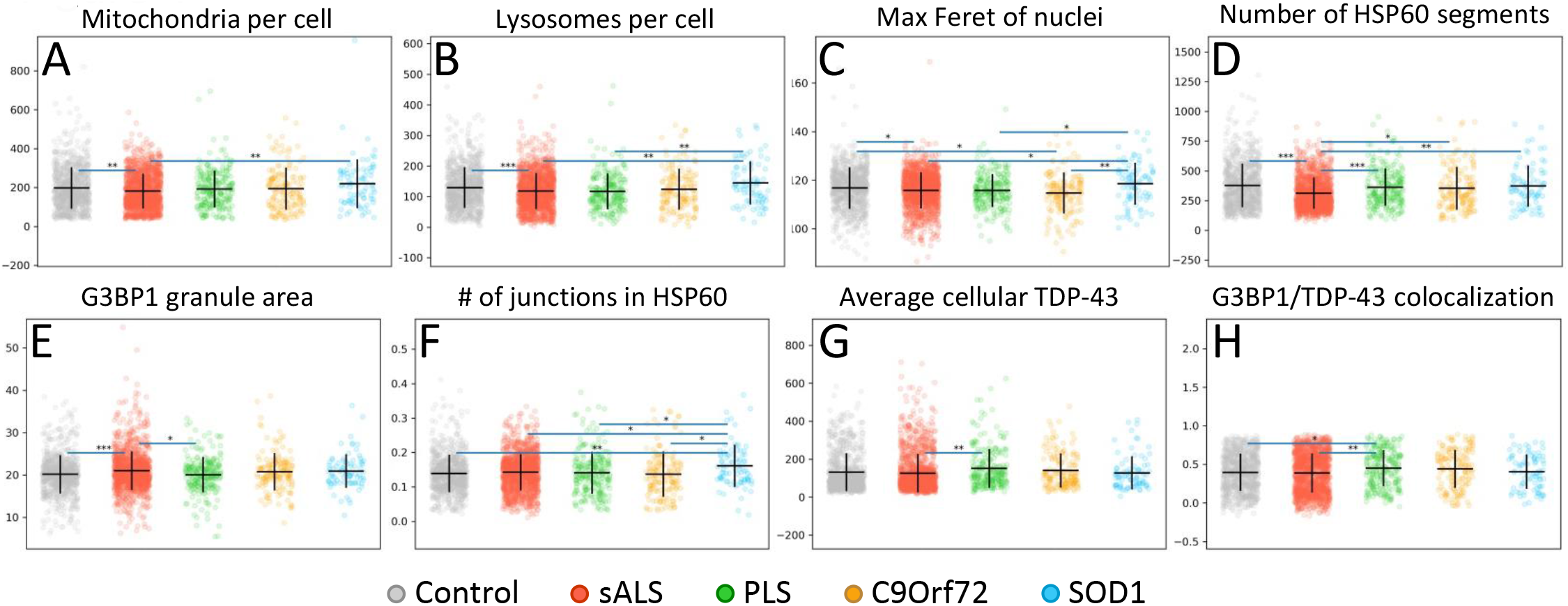
Examples of eight morphometric differences between disease groups (mitochondria per cell, lysosomes per cell, max feret of nuclei, number of HSP60 segments, G3BP1 granule area, number of junctons in HSP60, average TDP-43 intensity, TDP-43 and G3BP1 colocalization). Detailed description of features and measurement units can be found in Table S1. Each point represents a fibroblast line at a different perturbation (Control n=137×6, sALS n=225×6; PLS n=41×6; C9Orf72 n=26×6; SOD1 n=14×6). The complete lists of significant changes is shown in Fig. S2. Horizontal lines represent the mean; vertical lines represent the standard deviation; *: p-adj<0.05; **: p-adj<0.01; ***: p-adj<0.001.

Altogether, these findings highlight a few subtle measurable differences in morphometric features among control and the different ALS fibroblast groups. However, this data also indicate that individual morphometric features are clearly insufficient to serve as biomarkers to identify ALS groups and suggest that approaches that combine multiple features are needed.

### Dimensionality reduction shows clustering driven by perturbation but not by disease

We used dimensionality reduction to explore clustering, which approach was previously used with success by our group to identify a metabolically distinct subgroup of ALS(38). When projecting the morphometric features from LI and IF using t-distributed stochastic neighbour embedding (t-SNE) to two dimensions, the data from both imaging modalities showed separation driven by the different stress perturbations (Fig. 5A-B). On the other hand, no obvious clustering of disease groups was observed when using t-SNE (Fig. 5C-D). This analysis was also performed using only the data obtained under baseline conditions to prevent the dimensionality reduction algorithm to be driven by effects of the perturbations, but disease group clusters did not emerge (Fig. 5E-F). Using hierarchical clustering, which allows for clustering without compressing the data, also failed to find clusters of disease groups. Overall, these findings confirm that multivariate approaches can identify perturbations, but also that such unsupervised methods are inadequate to identify disease clusters based on morphometric features of ALS fibroblasts.

**Figure 5.**
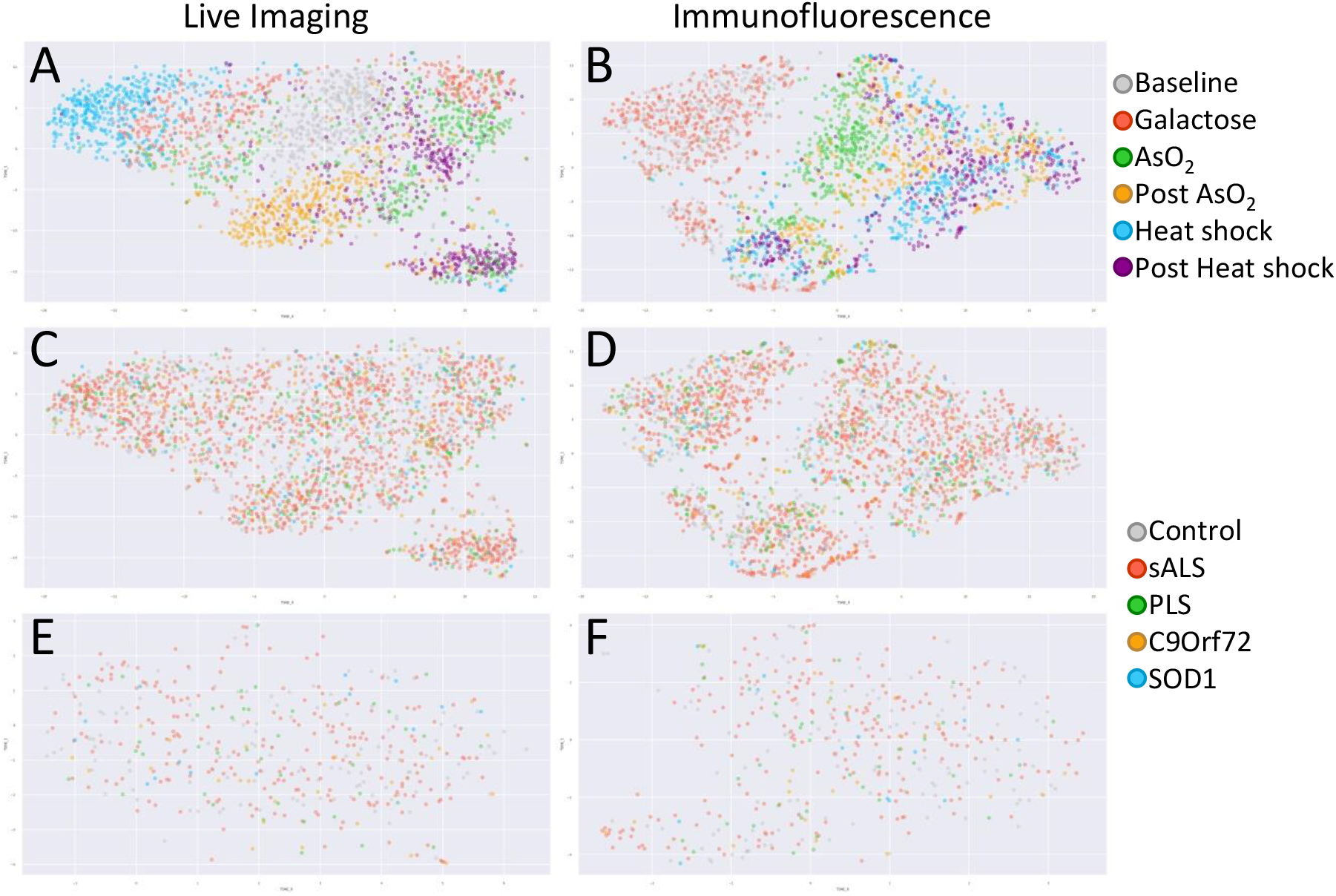
t-distributed stochastic neighbor embeddings. Projections of the two imaging modalities to 2 dimensions (A, C, E: Live imaging; B, D, F Immunofluorescence). In A-B, fibroblast lines are colored by perturbation. C-D show the same data as in A-B, but the fibroblast lines are colored by disease group. In E-F, only baseline conditions are shown and fibroblast lines are colored by disease group.

### Machine learning techniques classify stress perturbations but have moderate ability to predict disease groups and clinical phenotypes

We applied machine learning algorithms to classify cell lines based on stress perturbations, disease state, and patient features. As shown in Fig. 1, four ML strategies were used: 1) applying random forest models to the patient level morphometric features (Morph+RF); 2) applying the bag of visual words algorithm to the cell level morphometric features (BoVW+RF); 3) applying random forest models to feature embedding data generated from the images using a pre-trained ResNet50 deep learning model (Emb+RF); and 4) training a ResNet50 deep learning model from scratch (DL-scratch) with modified input dimensions to fit our 5 channel datasets. These four approaches differ most notably in feature extraction. The Morph+RF approach uses the morphometric features that were extracted (Fig 2). These features were ranked by the RF model in order of importance for classification for the given task. In the case of BoVW+RF, the same morphometric features were used, but this approach uses features generated from the distribution of the cell level measurements instead of using averaged patient level data. Unlike Morph+RF, this approach can determine if only a subset of cells of the patient hold valuable information for the classification. The Emb+RF approach uses transfer learning, taking advantage of automatically generated features that were learned by training on the ImageNet dataset. This approach could find features that we had not considered when designing our morphometric feature extraction. Finally, in the DL-scratch approach, features are automatically generated using our specific imaging dataset. This approach generates more relevant features for our data but takes much more computational work to train than the RF approaches.

We formulated nine classification tasks and divided the fibroblasts into training and test sets (4:1 respectively on the cell line level). Dividing into sets on the line level is important to accurately measure model performance, because ML techniques were shown to recognize individual fibroblasts of the same line(44), and we saw inflated scores when set divisions were made on the single cell level. The tasks and the number of examples in the total, training, and test sets, broken down by class, are summarized in Table 1. Assignment of fibroblast lines to training and test sets was kept consistent for each classification task among the four ML approaches and for the tasks to distinguish stress perturbations and disease groups (Control vs ALS; sALS vs other ALS; sALS vs PLS). We used 5-fold cross-validation for the DL-scratch models and 10-fold for the three RF approaches. The area under receiver operating characteristic curve (ROC-AUC) was calculated for the test set and for each cross-validation set, which was then used to calculate the mean and 95% confidence intervals.

**Table 1.**
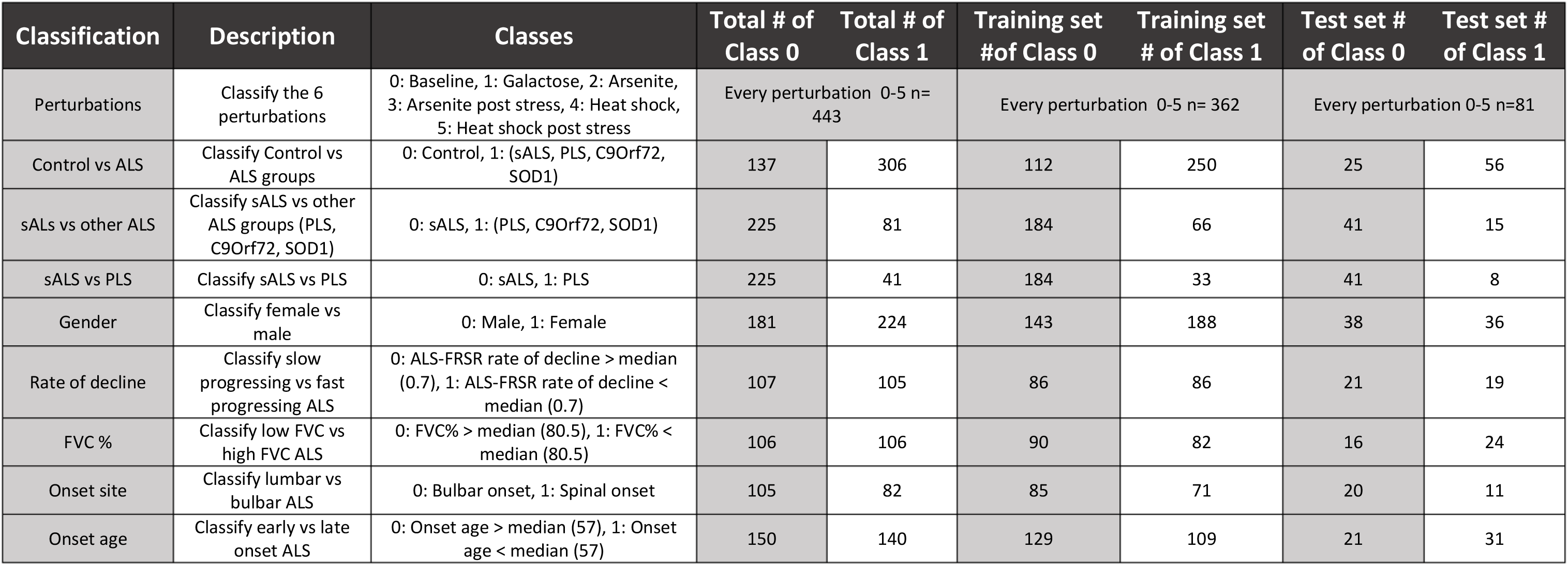
Detailed description of the classification tasks, class labels and numbers of examples in the training and test sets and the complete dataset.

| Classification | Description | Classes | Total # of Class 0 | Total # of Class 1 | Training set # of Class 0 | Training set # of Class 1 | Test set # of Class 0 | Test set # of Class 1 |
| --- | --- | --- | --- | --- | --- | --- | --- | --- |
| Perturbations | Classify the 6 perturbations | 0: Baseline, 1: Galactose, 2: Arsenite, 3: Arsenite post stress, 4: Heat shock, 5: Heat shock post stress | Every perturbation 0-5 n= 443 |  | Every perturbation 0-5 n= 362 |  | Every perturbation 0-5 n=81 |  |
| Control vs ALS | Classify Control vs ALS groups | 0: Control, 1: (sALS, PLS, C9orf72, SOD1) | 137 | 306 | 112 | 250 | 25 | 56 |
| sALS vs other ALS | Classify sALS vs other ALS groups (PLS, C9orf72, SOD1) | 0: sALS, 1: (PLS, C9orf72, SOD1) | 225 | 81 | 184 | 66 | 41 | 15 |
| sALS vs PLS | Classify sALS vs PLS | 0: sALS, 1: PLS | 225 | 41 | 184 | 33 | 41 | 8 |
| Gender | Classify female vs male | 0: Male, 1: Female | 181 | 224 | 143 | 188 | 38 | 36 |
| Rate of decline | Classify slow progressing vs fast progressing ALS | 0: ALS-FRSR rate of decline > median (0.7), 1: ALS-FRSR rate of decline < median (0.7) | 107 | 105 | 86 | 86 | 21 | 19 |
| FVC % | Classify low FVC vs high FVC ALS | 0: FVC% > median (80.5), 1: FVC% < median (80.5) | 106 | 106 | 90 | 82 | 16 | 24 |
| Onset site | Classify lumbar vs bulbar ALS | 0: Bulbar onset, 1: Spinal onset | 105 | 82 | 85 | 71 | 20 | 11 |
| Onset age | Classify early vs late onset ALS | 0: Onset age > median (57), 1: Onset age < median (57) | 150 | 140 | 129 | 109 | 21 | 31 |

We found that under baseline conditions, only models that used IF features produced cross-validation scores higher than 0.5 with 95% confidence in distinguishing disease groups (Table 2). The Morph+RF approach did so at the Control vs. ALS (0.62 ±(0.07)) and sALS vs. other ALS (0.63 ±(0.11)) tasks, while the BoVW+RF approach did so at all three disease group prediction tasks (Control vs. ALS: 0.63 ±(0.10), sALS vs. other ALS: 0.62 ±(0.09)), sALS vs. PLS: 0.60 ±(0.05)). The Morph+RF approach yielded a ranking of feature importance for classification, highlighting nuclear intensity, number of HSP60 objects, G3BP1 granule minimum feret size, and G3BP1 granule area as most relevant for the Control vs. ALS, and number of HSP60 objects, cellular area, number of G3BP1 granules, cellular minimum feret size, and TDP-43 cytosolic ratio for the sALS vs. other ALS tasks (Table S2). We also trained models to predict clinical parameters, including rate of decline in ALS functional rating scale revised (ALS-FRSR), respiratory forced vital capacity (FVC%) at time of biopsy, age at onset and onset site (lumbar or bulbar), and sex. Sex and onset site are binary, while for continuous variables (rate of decline, FVC%, age at onset) samples were divided into two equal groups (upper and lower quantiles) by the median value. In these tasks, only the DL-scratch approach on IF showed significant performance at distinguishing early vs late onset ALS (0.60 ± 0.07), however, the test score was outside of the cross-validation score range.

**Table 2.**
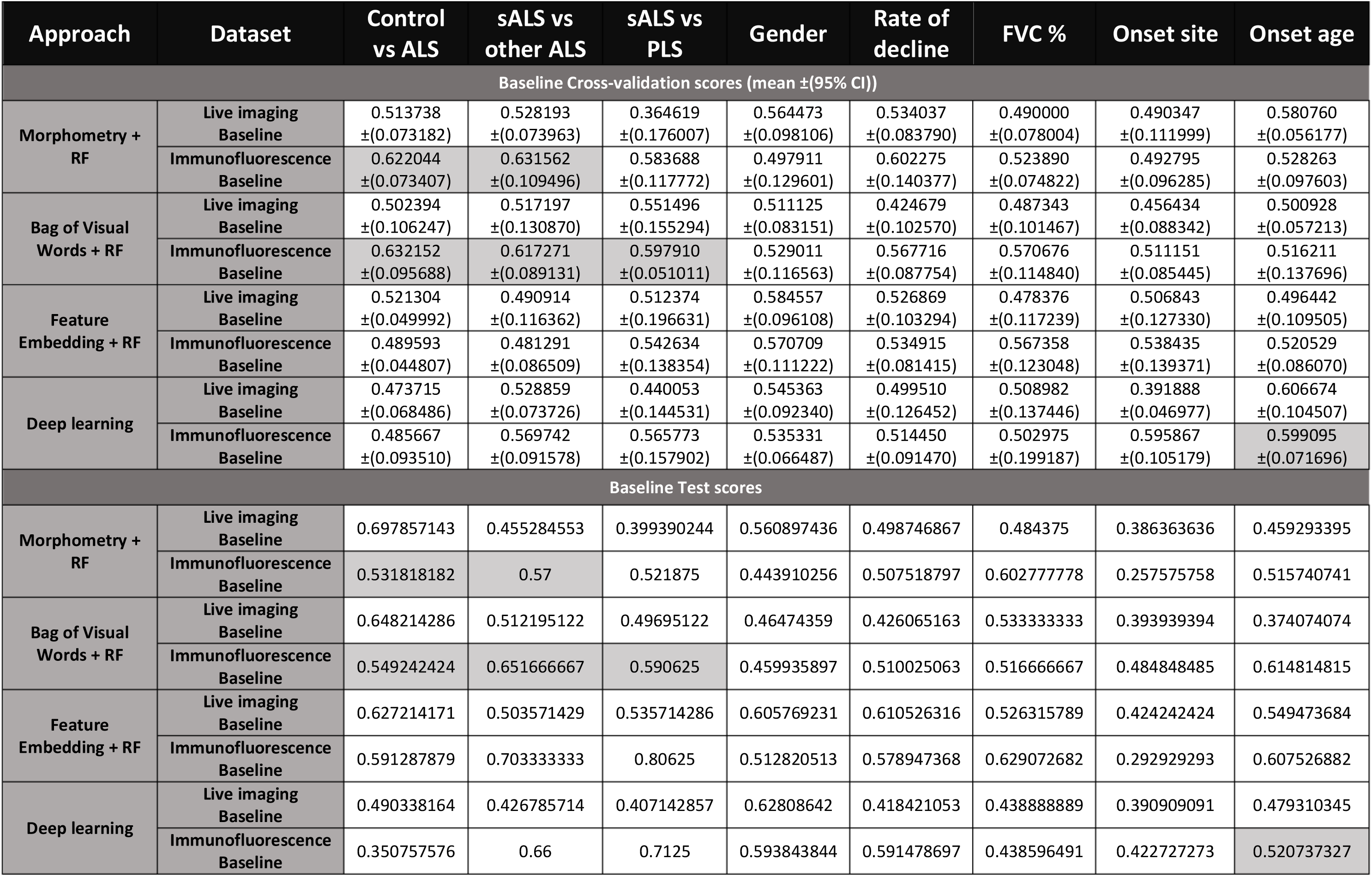
ROC-AUC values of the four ML approaches under baseline culture conditions for the different classification tasks for the cross-validation and test sets. Scores for models where the value is above 0.5 with 95% confidence on the cross-validation are marked by light gray for both sets.

Next, we explored whether perturbations could be predicted by ML approaches (Table 3). We tested the LI and IF datasets separately and generated an ensemble model by averaging the classification probability scores from both imaging modalities (LI+IF Ensemble). All models produced almost perfect scores on both cross-validation and test examples, and ensembles achieved slightly higher scores than the individual models in all four ML approaches, indicating that both imaging modalities have highly relevant information regarding the type of perturbation, and that they also have some complementary information. For the Morph+RF approach, ER tracker intensity, lysotracker intensity, and the number of mitochondrial junctions in LI, and G3BP1 granule number, TDP-34 nuclear intensity, and TDP-43 cytosolic ratio in IF were the most important features for predicting stress perturbations (Table S2).

**Table 3.**
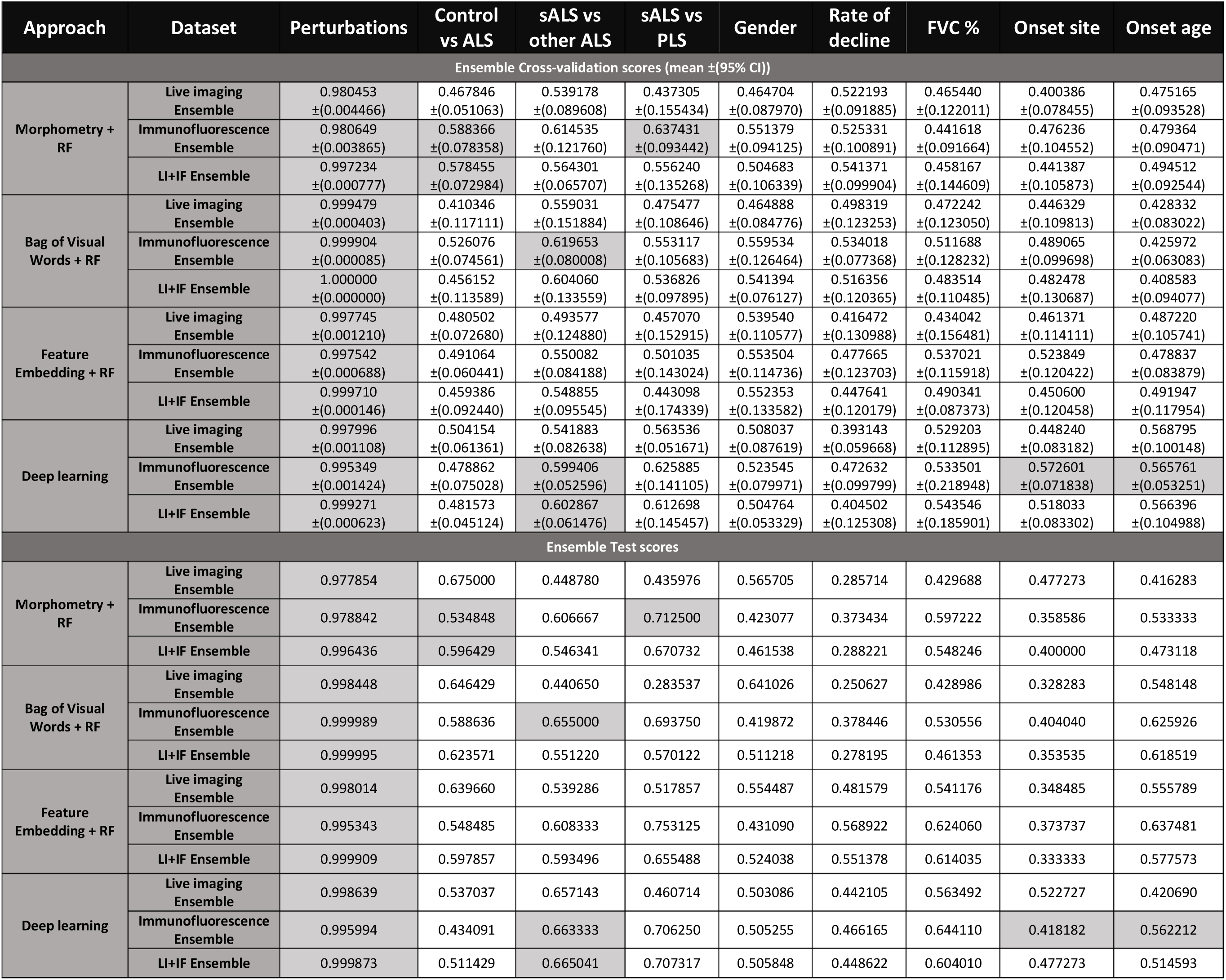
ROC-AUC values of the four ML approaches using ensembles for the different classification tasks for the cross-validation and test sets. Scores for models where the value is above 0.5 with 95% confidence on the cross-validation are marked by light gray for both sets.

Finally, we tested whether including stress perturbation data would increase performance scores on the classification tasks (Table 3). For each task, model building was done as for the baseline condition, yielding 12 classification probabilities per line (two imaging modalities x (baseline + five stress perturbations)). We then created three ensemble models by averaging the scores of the six models from each imaging modality or of all twelve models (IF Ensemble, LI Ensemble, IF+LI Ensemble). The Morph+RF approach produced ROC-AUC values that are higher than chance on the IF (0.59 ± 0.08) and LI+IF ensemble modes (0.58 ± 0.07) with 95% confidence on the cross-validation sets predicting Control vs. ALS. When predicting sALS vs. other ALS, we measured similarly modest but significant scores by the BoVW+RF IF ensemble (0.62 ± 0.08) and the DL-scratch IF (0.60 ± 0.05) and LI+IF (0.60 ± 0.06) ensemble approaches. When predicting sALS vs. PLS, only the Morph+RF approach on the IF ensemble (0.64 ± 0.09) was higher than random guessing. For predicting clinical parameters, the DL-scratch IF ensemble approaches showed scores barely higher than chance when predicting onset site (0.57 ± 0.07) and age at onset (0.57 ± 0.05), although the test score was well below 0.5 for the onset site prediction model (Table 3). Overall, the perturbation ensemble method scores showed success at the same task by the same approaches with little evidence of improvement from the baseline scores, and like under baseline conditions, we found no significantly scoring ML approach using the LI modality, indicating that the IF data was more valuable for these tasks than the LI. Our results suggest that the ML approaches that we developed based on fibroblast imaging can be highly successful in predicting stress perturbations but only modestly in predicting disease groups and disease phenotypes.

## Discussion

In an attempt to develop new biomarkers for ALS, we have tested a large collection of primary fibroblast lines in regular culture conditions and under different stress perturbations, combining live cell imaging of organellar morphology and function with immunocytochemistry of ALS-relevant proteins, such as TDP-43 and stress granule components. We performed morphometric feature extraction from the imaging data and found that stress perturbations, including oxidative stress, metabolic stress, and heat-shock, can be predicted with high accuracy using a variety of ML approaches. These results established proof of concept that the approaches that we developed are capable of learning from the data when the physiology of the cells is perturbed by strong environmental changes. Using the same parameters, we found some differences between control and ALS fibroblasts and between different ALS groups, but these differences had modest effect sizes and showed limited performance in predicting disease status. Therefore, the findings from this study suggest that organellar morphometry, mitochondrial membrane potential, TDP-43 localization, stress granule characteristics, and other potentially ALS-relevant features, both individually and in combination, under baseline or perturbed culture conditions, are not sufficiently indicative of disease state.

Our results partially agree with earlier reports on ALS fibroblasts. We previously reported increased energy metabolism in sALS and PLS(36, 37) fibroblasts, which we confirmed using metabolomics(38), but we and others did not observe hypermetabolism translating into mitochondrial morphology changes associated with the diagnosis of ALS(39, 45), highlighting the value of functional bioenergetic readouts over morphometry to detect metabolic changes in ALS fibroblasts. However, the present findings also partially diverge from previous reports. Several studies found abnormal TDP-43 expression in the skin of sALS patients by immunohistochemistry(20, 22, 24), as well as in cultured fibroblasts in both familial and sporadic forms of ALS, either under baseline or stress conditions(21, 23, 25, 26, 31). We only found modest increases in TDP-43 immunostaining derived morphometric features in PLS fibroblasts but not in sALS. Moreover, increased stress granule formation was observed under acute arsenite stress in FUS and C9Orf72 mutant fibroblasts^31^. However, with a similar stress protocol we did not find differences in a larger cohort of C9Orf72 mutant fibroblasts, although we did find increases in G3BP1 derived features in sALS vs. controls. There are several possible explanations to why these discrepancies exist in the nuances of sampling, experimental design, and data analysis. For example, some studies reported the number of cells in which TDP-43 mislocalization was observed, while we averaged cytosolic/nuclear intensity ratios for each cell. In this study we tested novel data analysis techniques such the BOVW approach, which takes histograms of the single cell level data as features, as well as deep learning approaches, which yield class probabilities at the single cell level. Both algorithms are sensitive to differences in cell subpopulations with mislocalized TDP-43 in the different groups. Despite the sensitivity of these approaches to single cell level data, we were unable to classify disease groups based on TDP-43 imaging.

ML methods for different fluorescence microscopy tasks have been developing rapidly, including segmentation, object detection, denoising(46), image focus scoring(47), in Silico fluorescent labelling(48-51), improvement of resolution(52), label free cell death detection(53), and diagnosis of neurodegenerative diseases(40, 44), in fibroblasts specifically. In this study, we used ML algorithms built on a large sample size and multiple features capable of capturing classes of environmental perturbations. However, we were unable to obtain promising results for clinical applications that could confidently distinguish ALS groups or even disease from control fibroblasts. These limitations could reside in the choice of biological readouts. We focused on a number of parameters that we could feasibly obtain, were logically associated with ALS, and previously employed to develop biomarkers in fibroblasts for SMA(40) and Parkinson’s disease(44). However, it could be necessary to test different sets of readouts, such as phosphorylated TDP-43(21), ubiquitination(23), or nuclear pore proteins that are altered in ALS(54). Additionally, different perturbations could be tested, including proteasome inhibition(23), autophagy arrest, or calcium overload. Moreover, high resolution microscopy techniques could be employed to detect more subtle differences between ALS groups and controls, although a previous study using a relatively small samples size with STORM-super resolution microscopy did not identify significant differences between controls and sALS(39). Finally, on the data analysis side, advanced ML techniques could be used to find relevant subpopulations of fibroblasts for the classification task, similar to the approaches currently used for cancer histopathology, such as multiple instance learning(55) or end-to-end part learning(56).

A limiting factor, directly linked to the hypothesis that fibroblasts can serve as a platform for biomarkers development with imaging approaches, may be related to the nature of the cell type. ALS fibroblasts may not express morphological differences sufficiently robust to be detected by our biomarker development pipeline. Therefore, fibroblast morphometry may be inadequate to yield high-quality biomarkers for ALS diagnosis, stratification, or surrogate endpoints. Patient derived cells that more closely recapitulate ALS disease phenotypes, including motor neuron and glia differentiated from induced pluripotent cells or directly from skin fibroblasts(57, 58) could provide alternative platforms for imaging biomarkers. Large scale efforts, such as the Answer ALS project(59) will likely be a valuable resource for imaging biomarker development in the future.

## Acknowledgements

This work was supported by Development Grant 602762 from the Muscular Dystrophy Association (to CK) and NIH/NINDS grant R35 NS122209 (to GM). This study used fibroblast samples from the NINDS Repository, as well as clinical data related to the samples. NINDS Repository sample numbers corresponding to the samples used are: (ND39022, ND39023, ND29422, ND29774, ND29149, AG13244, AG12598, AG13285, AG14284, AG11482, AG13150, AG13968, AG12657, AG12597, AG12851, AG11489, AG12964, AG12850, AG13144, AG13994, AG14048, AG13348, AG13990).

## Declaration of interest statement

The authors report there are no competing interests to declare.

## Data availability statement

Raw data and derived data supporting the findings of this study are available from the corresponding author CK on request.

## Notes

### Competing Interest Statement

The authors have declared no competing interest.

