## Supplementary Figures for "Machine learning approaches based on fibroblast morphometry confidently identify stress but have limited ability to predict ALS"

### Supplementary figure 1

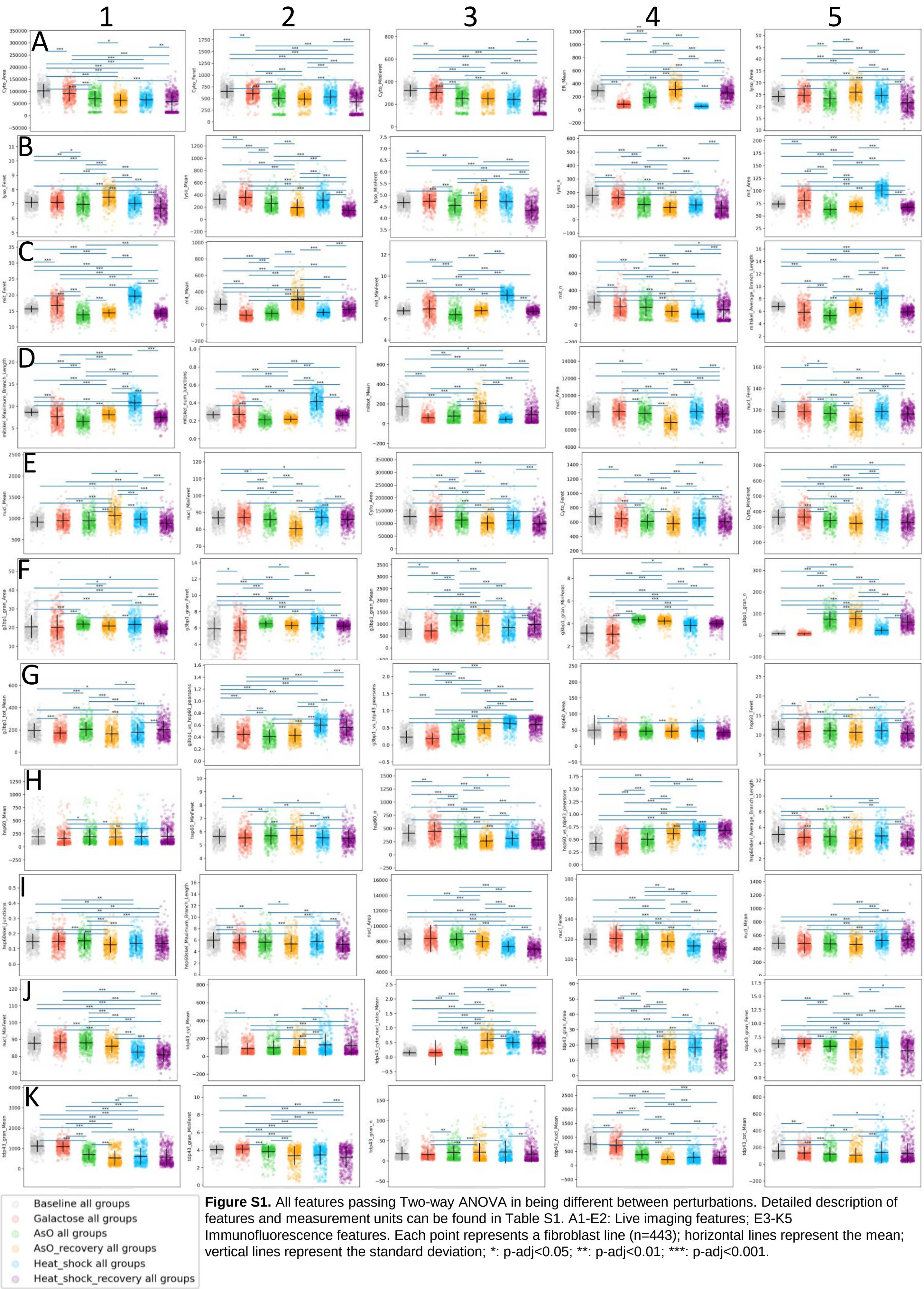


**Figure S1.** All features passing Two-way ANOVA in being different between perturbations. Detailed description of features and measurement units can be found in Table S1. A1-E2: Live imaging features; E3-K5 Immunofluorescence features. Each point represents a fibroblast line (n=443); horizontal lines represent the mean; vertical lines represent the standard deviation; *: p-adj<0.05; **: p-adj<0.01; ***: p-adj<0.001.

### Supplementary figure 2

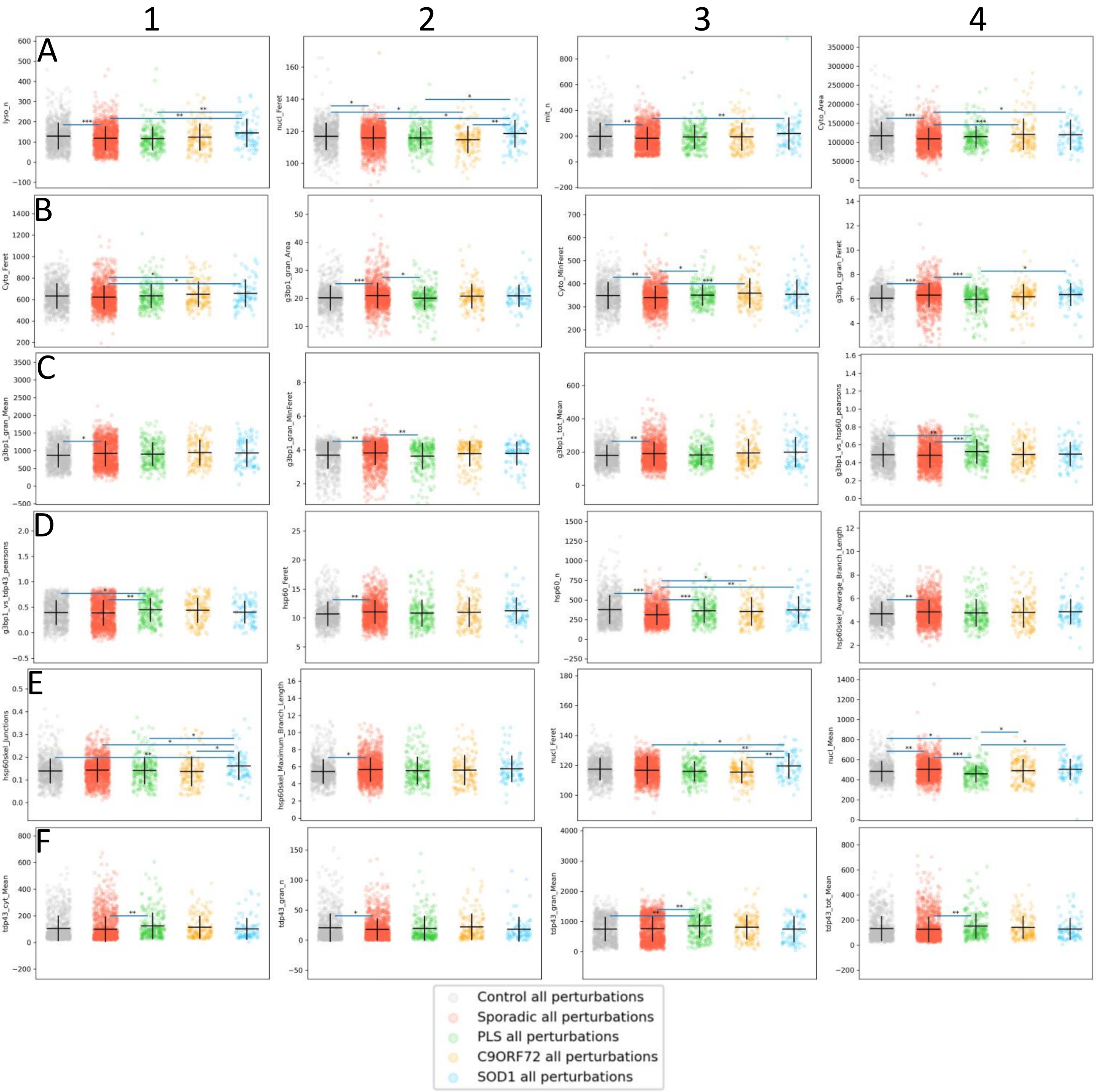


**Figure S2.** All features passing Two-way ANOVA in being different between disease groups. Detailed description of features and measurement units can be found in Table S1. A1-A3: Live imaging features; A4-F4 Immunofluorescence features. Each point represents a fibroblast line at a different perturbation (Control n=137x6, sALS n=225x6; PLS n=41x6; C9Orf72 n=26x6; SOD! N=14x6); horizontal lines represent the mean; vertical lines represent the standard deviation; *: p-adj<0.05; **: p-adj<0.01; ***: p-adj<0.001.

### Supplementary figure 3

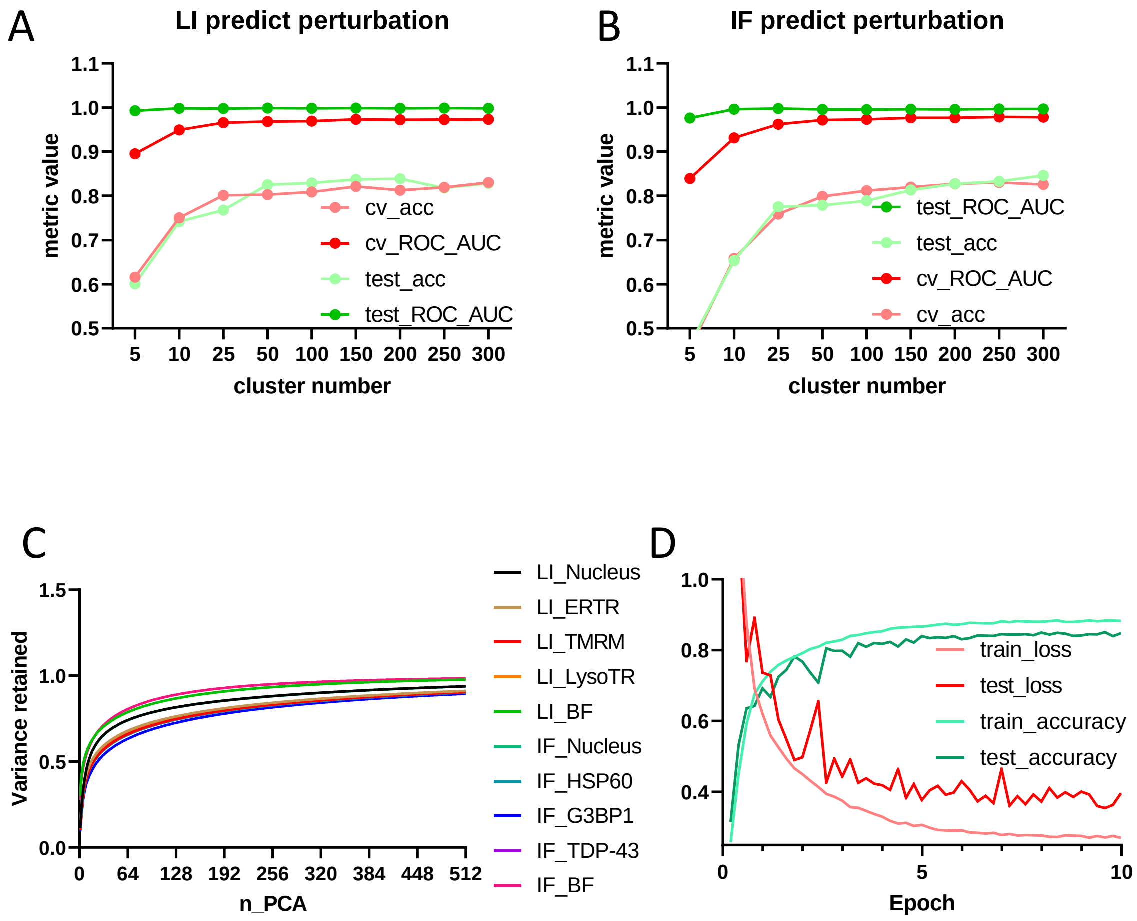


**Figure S3.** Model building quality control experiments. A-B test and cross validation accuracies and ROC-AUC values for the BOVW approach to classify perturbations using different k values for k-means on the live imaging(A) and Immunofluorescence (B) datasets. C: Variance retained by the PCA compression of the 2048 ResNet50 features for every channel in the two imaging modalities. D: Example training curves (deep learning approach to predict perturbations on immunofluorescence) throughout the 10 epochs of optimization.
